# The Lab Streaming Layer for Synchronized Multimodal Recording

**DOI:** 10.1101/2024.02.13.580071

**Authors:** Christian Kothe, Seyed Yahya Shirazi, Tristan Stenner, David Medine, Chadwick Boulay, Matthew I. Grivich, Fiorenzo Artoni, Tim Mullen, Arnaud Delorme, Scott Makeig

## Abstract

Accurately recording the interactions of humans or other organisms with their environment and other agents requires synchronized data access via multiple instruments, often running independently using different clocks. Active, hardware-mediated solutions are often infeasible or prohibitively costly to build and run across arbitrary collections of input systems. The Lab Streaming Layer (LSL) framework offers a software-based approach to synchronizing data streams based on per-sample time stamps and time synchronization across a common local area netowrk (LAN). Built from the ground up for neurophysiological applications and designed for reliability, LSL offers zero-configuration functionality and accounts for network delays and jitters, making connection recovery, offset correction, and jitter compensation possible. These features can ensure continuous, millisecond-precise data recording, even in the face of interruptions. In this paper, we present an overview of LSL architecture, core features, and performance in common experimental contexts. We also highlight practical considerations and known pitfalls when using LSL, including the need to take into account input device throughput delays that LSL cannot itself measure or correct. The LSL ecosystem has grown to support over 150 data acquisition device classes and to establish interoperability between client software written in several programming languages including C/C++, Python, MATLAB, Java, C#, JavaScript, Rust, and Julia. The resilience and versatility of LSL have made it a major data synchronization platform for multimodal human neurobehavioral recording, now supported by a wide range of software packages including major stimulus presentation tools, real-time analysis envirnoments, and brain-computer interface applications. Beyond basic science, research, and development, LSL has been used as a resilient and transparent back-end in deployment scenarios including interactive art installations, stage performances, and commercial products. In neurobehavioral studies and other neuroscience applications, LSL facilitates the complex task of capturing organismal dynamics and environmental changes occurring within and across multiple data streams on a common timeline.

## I. INTRODUCTION

Recording and modeling brain dynamics supporting active, natural cognition involving eye movements, motor and other behavior is becoming an integral part of neurobiological research and requires multimodal recording of an organism’s neural processes and interactions along with concomitant changes in its environment. Successful multimodal recording demands adequate temporal resolution and precise synchronization of concurrently recorded data streams. In human neuroscience, mobile brain/body imaging (MoBI) [Makeig et al., 2009] is a multimodal recording concept involving synchronized recording of brain, behavioral, and environmental data streams with near millisecond (msec) resolution. Maintaining synchronization at this scale between brain (electro/magnetoencephalography (EEG/MEG); functional near-infrared spectroscopy (fNIRS), etc.), behavioral (body motion capture and eye movement tracking), physiological (electromyography, EMG, etc.), and environmental data streams (video, treadmill, balance plate, robots, or other agent positions and forces, sensory stimulation, etc.) often requires multiple computer systems with no hardwired common clock to relate the timing of their outputs.

Here, we describe the Lab Streaming Layer (LSL), a software framework that is helping researchers across academic and industrial settings meet the challenge of multimodal recording through its ability to collect and synchronize data streaming from multiple devices and platforms connecting asynchronously to a local area network (LAN) with broad hardware and software compatibility. LSL is a freely available open-source project under the umbrella of a dedicated GitHub organization https://github.com/labstreaminglayer, plus individual core repositories available from the Swartz Center for Computational Neuroscience (SCCN) (meta-package and core library). A listing of over 150 known LSL-compatible device classes is compiled at https://labstreaminglayer.org, which also serves as a landing page to tooling, documentation, and other resources. LSL is supported by an active international community of contributors (including several coauthors). Currently, two annual workshops in Europe and the U.S., bring together platform users, contributors, and developers, and present learning opportunities for newcomers. Organizers currently include the SCCN and teams at the University of Oldenburg and TU Berlin. The popularity of LSL cannot be explained by any one of its features. Rather, its focus on ease of use and robustness, and its distributed model that allows synchronization of a wide mix of applications from multiple vendors and open-source projects running on multiple computers (desktop or mobile) contribute to its appeal, as does its broad platform compatibility with most major programming languages and all major desktop and mobile operating systems. The large LSL ecosystem and installed base also contributes to its growing adoption and appeal.

One of LSL’s technical features is the synchronization of distributed neuroscientific data streams based on a peer-to-peer protocol modeled after the Network Time Protocol (NTP) as specified in RFC 5905 [Martin et al., 2010]. A closely related component is LSL’s decomposition of timing error into three components: a constant, a slow-varying, and a noise component, which are each addressed separately. Using these two approaches, LSL can ensure that timestamps associated with every data sample, collected across multiple acquisition devices and computers, are accurately compensated for intrinsic device delay, clock drift, and jitter in the presence of variable network transmission latency. This capability is crucial in neuroscience research where near-msec precision can be essential for accurate data analysis and interpretation, particularly in studies involving complex brain/body dynamics, high-intensity biomechanics, and multi-subject interactions.

Challenges in collecting proper multimodal recordings include 1) the need to synchronize data streams from different platforms, 2) including data streams with heterogeneous sampling frequencies, 3) set up and staff training of multiple recording workstations and (possibly proprietary) software, 4) interfacing with multiple proprietary data access APIs with limited OS and programming language support, documentation, and learning resources, and 5) meeting challenges in data conversion, integration, storage, sharing, and reproducibility. Several hardware synchronization tools have been developed to address the pre-sampling synchronization in multimodal recordings. These include intricate systems of TTL (transistor-transistor logic) pulses, equipment for measuring throughput delays of recording instruments, and dedicating one instrument recording channel as a synchronizing clock [Artoni et al., 2017], [Maidhof et al., 2014], [Bannach et al., 2009].

Recent advances in hardware-managed synchronization can improve common clock accuracy for digitally triggered events to tens of microseconds, including solutions based on shared clocks and analog-to-digital (A/D) converters and [Chuang et al., 2021] radio-frequency trigger modules [Cerone et al., 2022]. However, the use of hardware data synchronization approaches is very often not feasible in laboratories without resources to engineer special-purpose solutions across the range of proprietary acquisition systems researchers wish to use in their experiments. This is still more the case for low-cost and/or consumer-grade microelectronics-based systems that can now be used to record multimodal data inexpensively in paradigms, allowing, among others, greater degrees of participant mobility or at-home use.

Heterogeneous sampling frequency, platform inaccuracies, jitter, and sampling fluctuations make synchronization of the data stream using ‘start/stop’ events insufficient for neuroscience purposes. Such a setup may cause synchronization to drift by many milliseconds within mere minutes of data collection, which typically grows longer over longer recording durations. A recent study of multimodal MoBI data collection methods concluded that frequent TTL pulses are needed to retain millisecond synchronization between data streams [Artoni et al., 2017]. Without this or some other hardware or software organizing method, data streams with different sampling frequencies typically drift out of synchronization over time, compromising their worth for joint analysis.

The setup and maintenance of professional timing equipment across multiple workstations running mutually incompatible recording software is time-consuming and may require a dedicated recording technician and/or experimenter team to run, monitor, and document the data collected by each system. A dedicated staff training process is often required to learn to operate the acquisition software associated with each system.

Finally, owing to the proprietary nature and variety of data collection software and data access means for different systems and the need to record metadata stored in different forms and locations, performing data conversion and preprocessing, integration, annotation, storage, analysis, and sharing is challenging. All these factors limit access to high-quality research capabilities.

### A. The Broader Landscape in Multimodal Recording

The LSL project was started in 2012 in response to an emergent need for robust multi-modal data acquisition at the Swartz Center for Computational Neuroscience (SCCN), UCSD, by the first author (Christian Kothe), where also the multimodal Mobile Brain/Body Imaging (MoBI) concept was originally proposed and first demonstrated [Makeig et al., 2009]. The available software at the time for this purpose was a partly proprietary package that was then in use at SCCN. Another technology predating LSL is the Tobi Interface A [Breitwieser and Eibel, 2011], which mainly aimed to standardize the representation of biosignals. HLA Evolved [Möller et al., 2008] was another solution for robust distributed simulator event tracking, which influenced our attention to reliability. There was no real-time data access protocol natively supported by multiple vendors of EEG hardware, let alone of a broader spectrum of neurobehavioral modalities.

Shortly after availability, LSL grew rapidly in popularity and found enthusiastic supporters both among academic labs and hardware manufacturers. As of mid-2025, LSL has been mentioned more than 2300 times in scientific articles, and is supported by the majority of popular real-time processing platforms for brain-and bio-signals, including BCI2000 [Schalk et al., 2004], OpenViBE [Renard et al., 2010], NeuroPype (Intheon, La Jolla, CA), Open Ephys ^1^, BCILAB [Kothe and Makeig, 2013], and MNE-Python [Gramfort et al., 2013], and younger platforms such as Timeflux [Clisson et al., 2019], MEDUSA [Santamaría-Vázquez et al., 2023], and Dareplane [Dold et al., 2023]. Since most high-level processing frameworks have a modular data source concept, most other brain-/bio-signal processing platforms can be made LSL compatible with relatively little effort and can thereby be made to leverage the full breadth of LSL-supported hardware. LSL has also been chosen as an underlying transmission protocol by commercial multi-vendor system integrators, including iMotions^2^ and BrainProducts^3^.

Since LSL is simultaneously a publish/subscribe overlay network and API, a time-synchronization solution, a multi-modal time-series and meta-data recording solution, and a real-time streaming tool with native support for event data, there are to our knowledge not many directly comparable alternatives. When reduced to its network protocol aspect, some alternatives are ZeroMQ^4^, MQTT^5^, plain TCP/IP, and Redis^6^ (e.g., as used in BRAND [Ali et al., 2023]). In the audio control domain an established protocol is Open Sound Control (OSC). Besides Open Epyhs, another project supporting multiple types of electrophysiology hardware is BrainFlow^7^, which currently supports a range of low-cost and DIY devices. For instrument and lighting control, respectively, well-known examples with good timing support are MIDI and DMX, but these do not leverage existing Ethernet or Wifi networking. However, it should be noted that even these solutions can, and some have been, integrated with LSL via bridge adapters. Alternatives for time synchronization are the precision time protocol (PTP) [IEEE SA Standards Board, 2020], which requires dedicated hardware, and manual NTP-based synchronization. Without a doubt, numerous research labs have developed countless pieces of in-house software that acquires data from two or more devices, some of which are also open-source projects (e.g., Bonsai [Lopes et al., 2015] with its focus on video and electrophysiology analysis of behaving rodents mainly on Windows workstations), but to our knowledge, none currently enjoy a degree of popularity, broad plug-and-play device compatibility, and large installed-base as LSL.

### B. LSL Limitations

Despite the stringent LSL time synchronization guardrails described below, LSL performance has some limitations. Most importantly, LSL does not have access to any incoming data *until* the moment it is received by the microprocessor (CPU) or microcontroller unit (MCU) on which the LSL software communicating with the device is running. Thus LSL cannot itself learn or estimate whatever *on-device delays* within each recording device occurred (the intervals accruing between data signal input and its arrival in the software). Measuring on-device delay (and ideally histogram) at least once for each acquisition stream is, therefore, necessary to allow LSL to convert the recorded times of data arrival into times of data capture. Once known, the delays, which LSL models as constant in between setup changes, can be accounted for and declared in software. This limitation is inherent to multimodal neuroscience data acquisition systems engineered without common hardware clock availability.

### C. LSL Advantages

The LSL approach to synchronized aggregation of concurrent data streams has three main advantages that together significantly enhance the data acquisition process: 1) Facilitating multi-modal data collection with heterogeneous and/or irregular sampling rates, 2) enabling distributed measurement and data processing across multiple systems, and 3) streamlining both real-time and offline access to time-stamped multimodal data through its companion *XDF* file format.

The LSL unified Application Programming Interface (API) and protocol standardize data exchange across any number of measurement modalities, creating a consistent real-time data stream access interface. This simplifies initial device setup, allowing LSL-compatible clients to require minimal or often no modifications to function with devices from different vendors. The API also offers the flexibility to use several of the most popular programming languages, allowing it to be integrated into almost any piece of existing software with little effort.

LSL allows time-synchronized stream readouts from all networked devices, simplifying the experimental process to merely starting the included recording devices and melding the received streams into an integrated XDF data record using the LSL *LabRecorder* application (or any equivalent of choice), eliminating the need to manage multiple data file formats and increasing the efficiency of either near-real time or *post hoc* data analysis. Moreover, LSL network protocol standardization facilitates the distribution of data measurement and processing across multiple computers without explicit network parameter configuration, increasing data acquisition versatility.

## II. SYSTEM OVERVIEW

LSL is a local network that runs on top of (or *overlays*) an Internet Protocol (IP) network running at the experiment site. LSL network peers can **publish** and **subscribe to** any number of **streams** of single-or multi-channel time-series data (Figure 1). LSL regularly quantifies clock offsets (OFS) and round-trip time (RTT) between peers to enable data stream synchronization. Multi-channel samples of any stream published on LSL contain the channel values (of flexible type) and a time stamp assigned by LSL or the LSL integration (“LSL App”) for the device. Peer access to LSL is set up using a dynamic library (*liblsl*) available for most POSIX-compatible platforms [IEEE SA Standards Board, 2018] including Windows, Linux, MacOS, Android, and iOS. The LSL API has been designed to “hide” the complexities of time synchronization and real-time network programming from both researchers and device manufacturers, while ensuring maximum network resiliency against dropped connections and data losses.

**Fig. 1.**
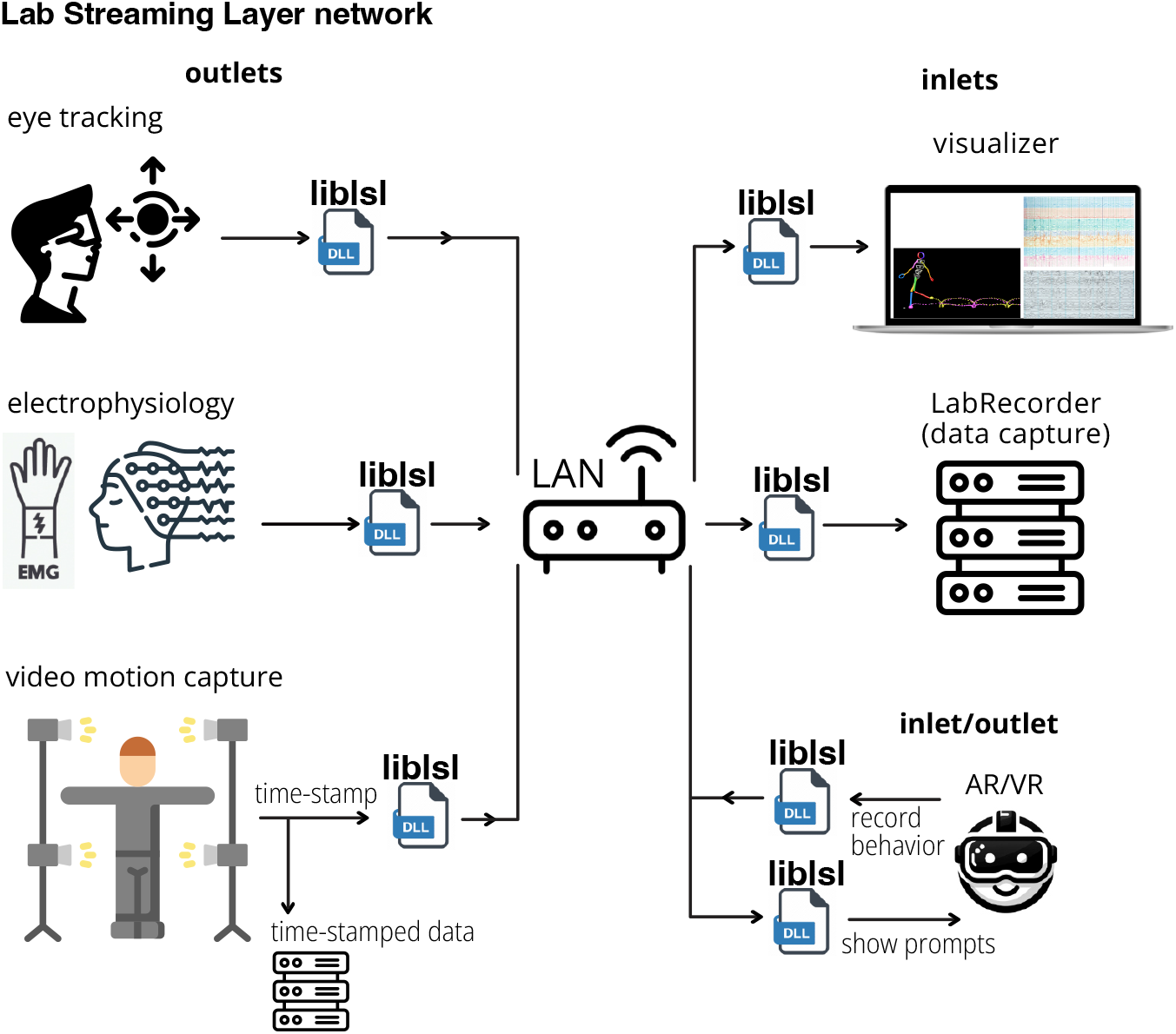
System overview. The Lab Streaming Layer (LSL) creates a *network* connecting data acquisition, storage, and processing devices overlaying the local network (LAN) on which they are streamed. LSL handles publishing and subscribing to data streams, clock synchronization, accounting for network delays, and jitter using the LSL dynamic library (*liblsl*). LSL *outlets* publish data streams to the network that LSL *inlets* can subscribe to. *LabRecorder* is a space-efficient and high-throughput LSL recording program that can supervise recording of streams from any number of LSL *outlets*. Clients on the network include device integrations (seen on the left-hand-side), single-or multi-stream visualization or real-time processing components, and arbitrary stimulus presentation and response collection mechanisms.

### A. LSL Objectives

Chief goals governing LSL construction were: **a)** to simplify the discovery and selection of the published streams, **b)** to simplify publishing of active data streams to subscriber applications in near real-time, **c)** to supply sufficient metadata to allow for full interpretation of the transmitted time series, **d)** to solve the time-synchronization problem for concurrent data streams with an error low enough for most neurobehavioral research (i.e., at most msec-scale), **e)** to provide adequate out-of-the-box fault tolerance across a range of commonly-encountered failure scenarios (such as single-device failures, reconnects, restarts, intermittent network connectivity loss, and so forth), **f)** to establish a unified multimodal data representation, and **g)** to offer an API to access, transmit, and (when needed) store data from any set of data streams, regardless of modality.

Other possible objectives were explicitly *not* LSL design goals: *a)* building an online or *post hoc* data processing system (although such systems can easily be built on top of LSL), *b)* building an internet-scale and/or internet-facing data transport system, *c)* replacing or competing with existing data acquisition software (e.g., device drivers or applications), *d)* replacing or competing with non-signal intra-process or inter-process message queuing systems, or *e)* solving needs far outside physiological or neurobehavioral research (e.g., high-energy physics).

### B. LSL Design

The LSL software framework consists of three main components: the LSL API and language wrappers, the LSL core library (*liblsl*), and the LSL protocols (See Figure 2).

**Fig. 2.**
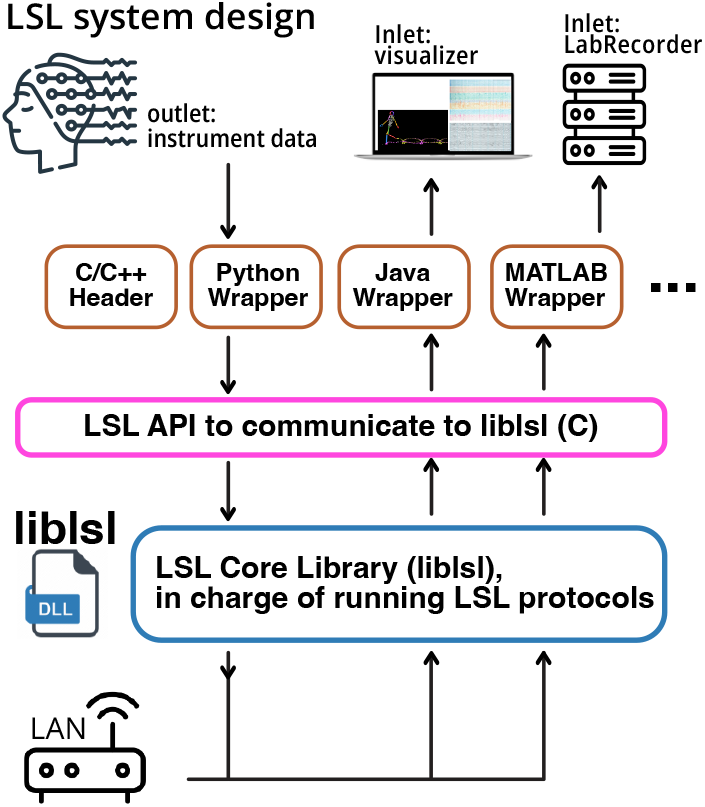
Lab Streaming Layer Design. LSL consists of three main components: **1** LSL language wrappers and API, **2** LSL core library (*liblsl* ), and **3** LSL protocols. The LSL API is a unified interface enabling communication with the LSL core library from external instruments and devices. The API was originally composed in C/C++ and is wrapped in other languages. The LSL core library (*liblsl* ) is written in C++ and implements all features that LSL offers. The LSL protocols are the set of steps and standards required to establish reliable communication and synchronization between peers.

**The LSL API** is a unified interface to communicate with the LSL core library from external instruments and devices. To maximize compatibility and ensure a stable Application Binary Interface (ABI), LSL presents a C API in agreement with shared-library best practices [Drepper, 2011], although the core is implemented in C++. Thanks to this stable ABI, support for other programming languages can be implemented with the C Foreign Function Interface (FFI), which enabled the creation of a wide range of wrappers for languages such as Java, C#, Python, Matlab, Rust, and several others. A header-only C++ API is also natively provided by the core library. These API wrappers provide the same metaphors, terminology, and functionality that the core C/C++ API provides. Since its initial release, liblsl has remained within the 1.x series, and all versions in this range are designed to be interoperable; connection handshakes negotiate the highest mutually supported protocol version to ensure compatibility. The library follows semantic versioning standards for API compatibility within major releases, while protocol versioning is handled separately; currently supporting two protocol versions that enable communication between different liblsl versions, including potentially decade-old software installations that remain critical in research environments.

Each existing API attempts to respect the idioms and standards of the language in which they are implemented. So, the Python API aims to be “Pythonic,” while the C API is an example of a “classical” C style, yet at the same time, all APIs cover an equivalent feature set. Developers can use the API to design executable programs to communicate with their peers on the network, publish data, and subscribe to streams from other peers.

A simple yet runnable example in Python that discovers, subscribes to, and then reads from an EEG stream on the LSL network is given in the following listing (equivalent examples are provided for all supported programming languages):

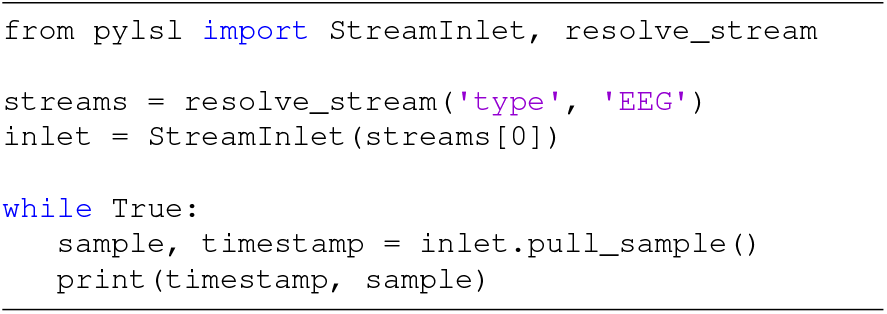

A corresponding simple example that generates 8 channnels of random floating-point numbers and streams them to LSL at approx. 200 Hz, here written in C++, is shown below. For best interoperability it is recommended to additionally specify meta-data such as channel labels, which is not shown here. Equivalent functionality is available for all other supported programming languages.

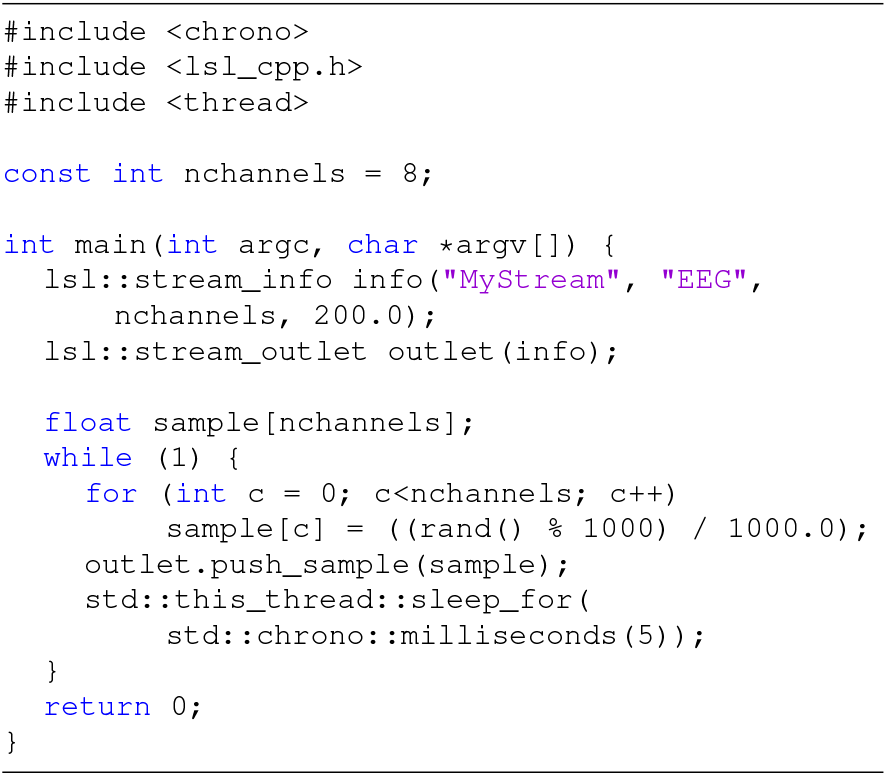

**The LSL core library** (*liblsl*) is written in modern C++ and^111^ manages features that LSL offers. Each peer needs to have a copy of the *liblsl* to communicate with other peers on the network. Our effort has been to maintain *liblsl* as a self-contained package to minimize its dependencies on packages that are not shipped with the LSL source code. Therefore, users should be able to compile the library in case compiled binaries are not available on a given platform.

Internally, *liblsl* uses *pugixml* [Kapoulkine, ] for XML and XPath processing, *loguru* [Delgan, ] for logging with configurable verbosity and log targets, and *Boost ASIO* [Kohlhoff, ], [Koranne, 2011] for portable high-performance asynchronous networking.

### LSL Network Protocols

LSL internally implements five network protocols to allow peers to create and maintain outlets to publish data streams, inlets to subscribe to streams, and to stream information objects each carrying all the requisite metadata for a data stream. By protocols, we mean the steps and standards to establish outlets, inlets, and metadata transfers. The five protocols are titled *(1)* Discovery, *(2)* Subscription, *(3)* Stream transmission, *(4)* Metadata transmission, and 5) Time synchronization. Adherence to the protocols is guaranteed by the core library (*liblsl*).

1. *) The Discovery Protocol:* The first stage in establishing communication between inlets and outlets is stream discovery. An application may discover outlet peers by broadcasting query messages into the network via UDP broadcast and UDP multicast (RFC1112) [Deering, 1989] to user-configurable multicast groups and awaiting responses. The query message contains an XPath 1.0 ^8^ compliant query string that specifies some metadata properties of the stream of interest (e.g., type=“EEG”). The host of each published stream on the network will then respond to matching queries with a small response packet that contains the essential properties necessary for establishing a connection specific to the querying peer so that a single machine can stream data to multiple peers at once. These include the name, type, and unique identifier of the stream and are formatted as an XML string. Responses to identical queries are cached for efficiency.

For convenience, all of this happens ‘under the hood’ of a single LSL function call. The programmer of an LSL application need not be concerned with the details of interfacing with a network stack for this to work. Furthermore, queries can be transported over several network protocols, including UDP broadcast and multicast of various scopes, and can be done using IPv4 and/or IPv6. LSL will correctly choose the right communication technique so that the programmer can be agnostic of the underlying network protocols.

The same LSL query protocol is used to automatically reconnect to a peer should the connection be lost during a data transfer – for example, if a software or network computer crashes, or a change in network topology occurs. Connection recovery will be successful even if the peer’s IP address has changed. This provides substantially greater resilience than most protocols that cannot recover from a change in IP addresses.

*2) The Subscription Protocol:* After a desired active outlet object is discovered, the host application on the subscriber side will want to connect a stream inlet to the outlet. This process is called an LSL subscription, enacted by establishing a TCP connection to a network endpoint advertised in response to the discovery query. A brief two-way protocol negotiation handshake establishes this connection. The handshake resembles HTTP/1.1 GET and its response [Fielding et al., ]. The purpose of this handshake is to exchange several transmission parameters such as the protocol version, byte order, buffer sizes, support for floating-point subnormals, etc.

A mutually agreed-upon sequence of test-pattern data is also transmitted to confirm that both parties can support the same protocol. The metadata header (stream information object) is also transferred from the host (outlet) to the client (inlet) to confirm that the endpoint does carry the requested data stream. Once this exchange is completed, the connection is formed, and time-series data will flow from the outlet to the inlet until the connection is terminated.

3) *The Stream Transmission Protocol:* LSL transmits time-series data as a byte stream split into packets by the underlying network layer. Samples in the time series may be marked for immediate transmission to enable use in real-time applications. This effectively indicates a ‘flush’ operation wherein the marked sample(s) are to be transmitted as soon as the underlying network permits. The byte stream is a sequence of encoded message frames. Every frame corresponds to one sample and includes a losslessly delta-compressed timestamp followed by the sequence of data values (bytes) encoded according to the format agreed upon during the connection handshake. While the underlying protocol is sample-oriented, the choice between immediate or deferred transmission allows users to send or receive time series either sample-by-sample or at the granularity of multi-sample chunks, where either side can choose to use either protocol, using easy-to-use high-level functions (the above code listing shows sample-wise sending and receiving).

4) *The Metadata Transmission Protocol:* In addition to time-series data, a stream’s metadata must be transferred from peer to peer. This metadata plays the same role as a file header in a time-series recording and contains information such as the stream name, type, channel count, sampling rate, etc. The metadata needs only be transmitted once and is thus treated by LSL as ‘out-of-band’ data. It is only transmitted on client request over a TCP connection. A simple connection handshake also precedes this transfer.

The metadata is plaintext and structured in accordance with an attribute-free subset of XML and can be of any length. LSL does not prescribe the metadata structure, but for interoperability, it is strongly recommended to adhere to a specification of content types (modalities such as EEG, Audio, Gaze, and so forth) and content type-specific nomenclature of XML fields. The type-specific nomenclature was co-developed with the XDF (extensible data format) project and is available online from the XDF GitHub Wiki. Since this metadata specification is plaintext XML, applications may extend and augment this metadata in any way that is suitable for a given data stream without breaking compatibility or deviating when necessary.

5) *Time Synchronization Protocol:* A common use case of LSL is streaming multimodal time series data from multiple peers to a separate peer that subscribes to (monitors and/or records) the multimodal data. LSL’s timestamping function returns the time of the most steady (i.e., monotonically increasing) high-precision computer clock available that has a minimum resolution of 1 msec or better (typically the machine uptime). The time offset between multiple computers’ clocks, as well as their relative drift, is continually measured and accounted for by LSL when synchronization information is utilized. When an inlet peer wishes to synchronize its clock with the respective outlet peer, a structured packet exchange is initiated following the basic NTP model. Since clocks need to be periodically re-synchronized due to the drift, this process will be repeated regularly (e.g., by default, every 5 seconds). LSL employs the clock filter algorithm of the Network Time Protocol (NTP) [Martin et al., 2010] to account for random spikes in network transmission delay. This process uses multiple packet exchanges to estimate the clock offset (OFS) and round-trip times (RTT) between peers in rapid succession (e.g., ten times across 200ms), yielding a set of OFSs and RTTs from which the one with the lowest RTT is retained.

Each packet exchange attempt for clock synchronization consists of a packet sent from the initiating peer to the receiver. This carries the local timestamp of the initiating peer and is noted as *t*_0_. The receiver then responds with two more timestamps, the receiving time of the original packet *t*_1_, and the time of resend *t*_2_. Upon receipt of this packet by the initiating peer, a final timestamp *t*_3_ is taken. Then,

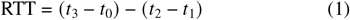

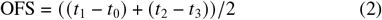

Therefore, RTT is the duration of the entire round trip minus the time spent on the receiving peer, and OFS is the averaged clock offset between the peers with symmetric network transmission delays canceled out. This measurement is a minimum-noise realization (because we choose the OFS at the minimum RTT) of the unbiased clock offset between the two peers. There can be a transmission time asymmetry between the forward and backward network path (e.g., due to driver implementation details), but the residual error after clock filtering is upper-bounded by the lowest delay of a machine’s network implementation and is therefore assumed to be well under 1 ms with most network hardware.

Using this time-varying measurement, the receiving side of LSL then constructs a model of the observed time stamps *t*_obs_ as a function of the time *t*_actual_ when the on-device measurement actually occurred (ignoring relativistic effects), an optionally smoothed estimate of the clock offset 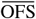, a device-specific constant offset *τ*, and a zero-mean noise term *ε*:

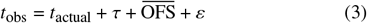

Using this formula, it is possible to recover *t*_actual_ for regularly sampled time series either using a recursive least-squares estimator in real time or linear regression in post-hoc data analysis, both of which are supported by LSL for the former and by XDF implementations for the latter.

### C. The Extensible Data Format (XDF)

The Extensible Data Format (XDF) is an open-source and general-purpose natively multi-modal container format for multichannel time series data with extensive associated metadata. XDF is tailored towards biosignal data such as ExG, GSR, and MEG, but it can also handle data with a high sampling rate (like audio) or data with a high number of channels (like fMRI or raw video). In general, every data stream collected by the LabRecorder, along with metadata and synchronization information is recorded into a single XDF file. Crucially, XDF follows the policy of recording all timing-related ground truth “as it happened”, which allows for post-hoc analysis and recovery of data in case of misbehaving devices or intermittent failures during a recording. A result of this choice is that, while XDF importers present a simple interface similar to that of many other file importers, XDF files represent an exact record of what occurred during an experiment, which can at times be complex, including a device disappearing and later (e.g., after an unplanned battery swap) reappearing.

In case of a high-bandwidth time series that may not be transferable over the network (such as uncompressed video), each frame of the stream may be timestamped and stored in the local machine (outlet) while the timestamp information and the metadata would be sent over LSL to the inlet machine and would be added to the XDF files. Another scenario in which this may be favorable is when video data falls under stricter privacy and regulatory requirements as personally identifiable information (PII) than most other information that can be recorded into an XDF file.

The XDF metadata is stored as XML content in an efficient binary chunk-oriented container file format, and the recognized metadata parameters are available at the XDF GitHub repository. XDF predefines an extensible set of content types (e.g., EEG, Audio, NIRS, and so forth) and associated metadata specifications, following a lightweight open process by which this specification is extended. This allows a single file to maintain comprehensive yet extensible modality-specific metadata on par with most unimodal biosignal file formats. XDF tools are available for download via the XDF GitHub page. A derived ANSI standard (ANSI/CTA-2060-2017) specifying a file format for a consumer-grade variant of XDF has since been published [ANSI, 2017].

### D. Failure Resilience

Preventing data loss is a major objective during data collection, especially in multimodal data acquisition where the probability of hardware issues grows linearly with the number of devices involved in a given data collection setup. LSL is equipped with a number of mechanisms for preventing catastrophic crashes and loss of data to ensure smooth operation, even in the event of computer crashes and lost network connections. To prevent data loss, LSL *outlet* and *inlet* objects can use variable-size buffers that have a configurable, arbitrarily large capacity. So, in case an *inlet* temporarily could not receive data from an *outlet*, the data can be buffered until the *inlet* can handle the transfer. The upper limit of all of this is the computer resources and network throughput.

In the event of an *outlet* dropping out, any *inlet*s connected to the *outlet* will attempt to reconnect. An event will trigger within the *inlet* to periodically search for the *outlet* and attempt to reconnect as soon as the *outlet* is rediscovered. Since the *outlet*’s information object can be created with a unique ID, this discovery will happen automatically even if the *outlet* is recreated on a different computer in the network and with a different IP address.

If an *outlet* drops out while an *inlet* is recording data, LSL can tolerate a discontinuity in the clock offset for the dropped stream after the rediscovery of the *outlet*, so that the outlet timestamp is consistent with the timestamp information prior to the dropout. This behavior is agnostic to the crash type and could resume recording of the discovered *outlet* even if the disconnection is a result of changing network topology, a computer crash and restart, or hardware failure like a dead battery.

Since these recovery processes happen automatically, the LSL user is shielded from having to cope with anything other than potentially a gap in a recorded data stream in the event that a device was intermittently not recording data. XDF tools typically come with built-in support for the detection and correct handling of such data gaps. These collective built-in efforts to recover connections between peers realize LSL’s failure resilience. While our validation tests focus on ideal conditions, LSL has been stress-tested under various failure scenarios—including device restarts, network congestion, and clock drift—to verify its resilience. Built-in mechanisms such as automatic reconnection, time offset renegotiation, and buffering help maintain data continuity under typical disruptions.

### E. Software Stack

LSL includes an ecosystem of applications to publish and subscribe to data streams, APIs in various languages built around the core dynamic library (*liblsl*), an extensible data recording format, *XDF*, post-hoc analysis for loading LSL synchronization performance, and tools for performing offline time-synchronization. This ecosystem can be accessed via the landing page and GitHub organization and meta-repository. LSL also offers rich and open-source documentation maintained by its developer community, available at https://labstreaminglayer.readthedocs.io.

However, it is far beyond the scope of this article to do justice to the greater LSL software ecosystem, which includes over a hundred compatible client applications, some open source and others vendor-native. Many applications in this greater ecosystem are hosted under an umbrella GitHub organization, while many others are vendor-provided data acquisition software with built-in LSL support, and an unknown number of further LSL clients can be found via internet searches. While this article focuses on acquisition devices, it is important to note that the LSL ecosystem also includes a robust collection of compatible stimulus presentation software, including most major programs used for this purpose, which are indispensable for scientific experimentation. Furthermore, the ecosystem includes software for real-time processing of collected data (for example, for brain-computer interface or neurofeedback applications), visualization, troubleshooting, experiment management, and various other tasks.

### F. Continued Development and Maintenance

Researchers and programmers from both academic and commercial sectors all over the world have contributed to the LSL source code and APIs. However, changes to the core library (usually bug fixes) are made very infrequently and with ultimate caution. Backward compatibility with existing applications is maintained at all costs. The bug rate is very low (less than one discovered every 6 months) and, so far, all bugs that were discovered were non-critical. Some bugs seen so far include a few memory leaks and typing errors in printing metadata and error messages. We have not found any bug affecting the proper operation of sending and receiving data (the primary LSL objective) in the past several years. Bugs in the LSL application ecosystem and APIs are more common, but given the stability and reliability of the core library and the simplicity of its interface, these bugs are relatively trivial to identify and cannot affect (i.e., crash) other LSL *inlet*s and *outlet*s – one of the less obvious benefits of a decentralized design.

To maintain stability, unit tests covering a wide array of both internal and API functions are run on all computing platforms for every change committed to the source code. In addition, the library is periodically stress-tested with hundreds of streams, randomized disconnects, shutdowns, reconnects, and randomized stream parameters. During such extreme network stress tests, some consumer-grade network equipment has been found to be less reliable (i.e., crashing) than the LSL implementation itself. Our dedicated benchmarks ensure that changes in operating systems and libraries do not impair the data exchange and synchronization performance. In addition, downstream libraries, such as mne-lsl, also follow continuous integration and unit testing best practices, providing additional implicit validation and stress testing of the LSL ecosystem.

## III. TESTING AND RESULTS

LSL has been extensively tested and validated by the biosignal research community in several studies [Bustamante et al., 2021], [Kang and Wallraven, 2023], [Weber et al., 2021], [Levitt et al., 2022], [Iwama et al., 2022], [Merino-Monge et al., 2020], [Blum et al., 2021], [Chuang et al., 2021]. Here, we provide some data concerning LSL’s performance on a local network (i.e., all LSL *inlet*s and *outlet*s running on a single machine), on a distributed network, and on a local network collecting data from mutiple instruments. We provide a simple yet effective recipe to determine, for a given data instrument, the total delay of the data path for a given instrument, which is a sum of the internal hardware delay (e.g., on-device buffers), wireless transmission latency and operating system, device driver, and driver access latency, which we term in the following the “setup offset” *τ*.

Using a scientific grade analog-to-digital/digital-to-analog I/O device (National Instruments Data Acquisition Box, NI-Daq, Austin, TX) we created a periodic pulse signal (Figure 3). We used the same NI-Daq to receive the same signal (<u>DataIn</u>), and create an <u>DataIn marker</u> when the pulse was going high. To create the DataIn marker, we chose the time the recorded signal reaches halfway to its maximum amplitude. We also recorded the <u>pulse event</u> directly from NI-DAQ using LSL.

**Fig. 3.**
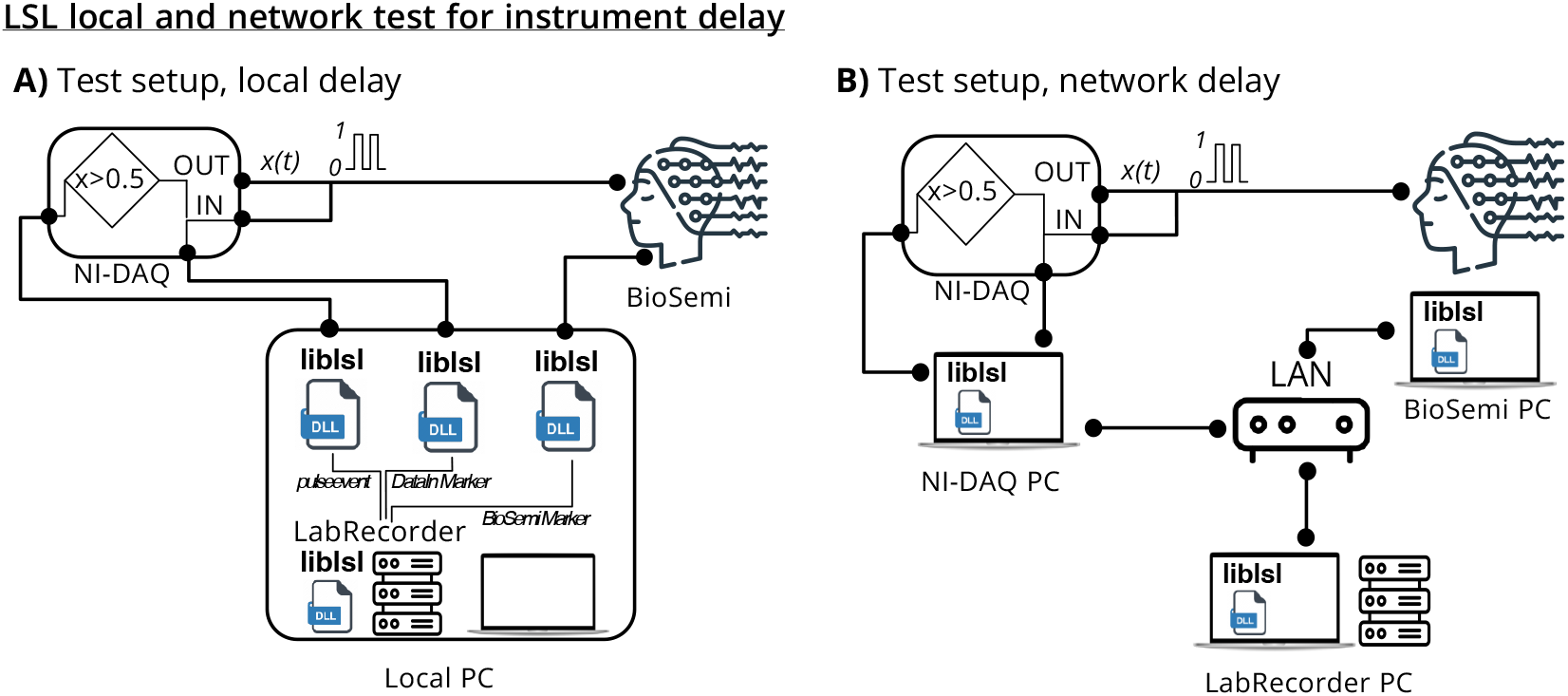
Synchronization performance setup. The setup consists of a National Instruments Data Acquisition Box (NI-Daq) that generates a periodic pulse signal (DataOut) and receives the same signal (DataIn). The same NI-Daq is used to create an LSL marker when the pulse is going high. At the same time, a BioSemi Active-II receives the same pulse signal as an LSL stream. The BioSemi stream and the marker stream are recorded using LabRecorder, the native LSL recording program. The LSL marker stream is used to calculate the synchronization accuracy of the BioSemi stream. **A**. The local setup is using a single computer to connect to the NI-Daq and BioSemi devices and record the streams using LSL LabRecorder. **B**. The network setup uses separate computers to connect to the NI-Daq, BioSemi, and the LSL LabRecorder.

At the same time, we used another scientific-grade signal recording device (BioSemi Active-II, BioSemi B.V., Amsterdam, the Netherlands) and read the same pulse signal as an LSL stream. We used a similar threshold for the BioSemi-recorded pulse signal (i.e., halfway to maximum amplitude, <u>BioSemi Marker</u>), so that we could add time markers when the pulse signal went high. We recorded the BioSemi stream and the LSL marker stream using *LabRecorder*, the native LSL recording program.

Finally, we compared the timestamps of the marker stream and the ‘high’ points of the BioSemi stream. The NI-Daq data input stream was sampled at 10 kHz, and the BioSemi data stream was sampled at 2048 Hz.

We expected to observe a constant offset (<u>setup offset</u>) between the two markers (i.e., DataIn Marker and BioSemi Marker) due to the setup and network topology, plus some jitter. We ran the NI-Daq controller, BioSemi, and LabRecorder on (1) a single machine (Intel Windows 7) to test the LSL’s local performance and (2) used separate network-attached machines for each of the NI-Daq controller, BioSemi, and LabRecorder (Intel Windows 7 for NI-Daq and Intel Windows 10 for each BioSemi and LabRecorder) to test LSL’s network performance. We analyzed the difference of 1500 high-points generated by NI-Daq and BioSemi systems to quantify jitter and setup offset.

Here, we purposefully avoided using state-of-the-art machines in order to test LSL performance on a more typical PC data acquisition setup.

### A. Instrument Latency in a Local LSL Setup

The results showed a five-microsecond lead time between the time a DataIn Marker was issued and the *pulse events* satisfied our defined threshold (Figure 4A). This is well below the 100-microsecond resolution of the NI-Daq reader, so we considered this lead time negligible. Comparing the BioSemi Marker and the DataIn Marker latencies indicated a 12.20 ms setup offset between the two markers (Figure 4B). The jitter of this offset (i.e., the standard deviation of the lag (see Figure 4B) was 156 microseconds, below the ∼500-microsecond Biosemi time resolution. Thus, the two streams could be aligned by removing this (pre-measured) device setup offset, and time jitter should not affect this alignment.

**Fig. 4.**
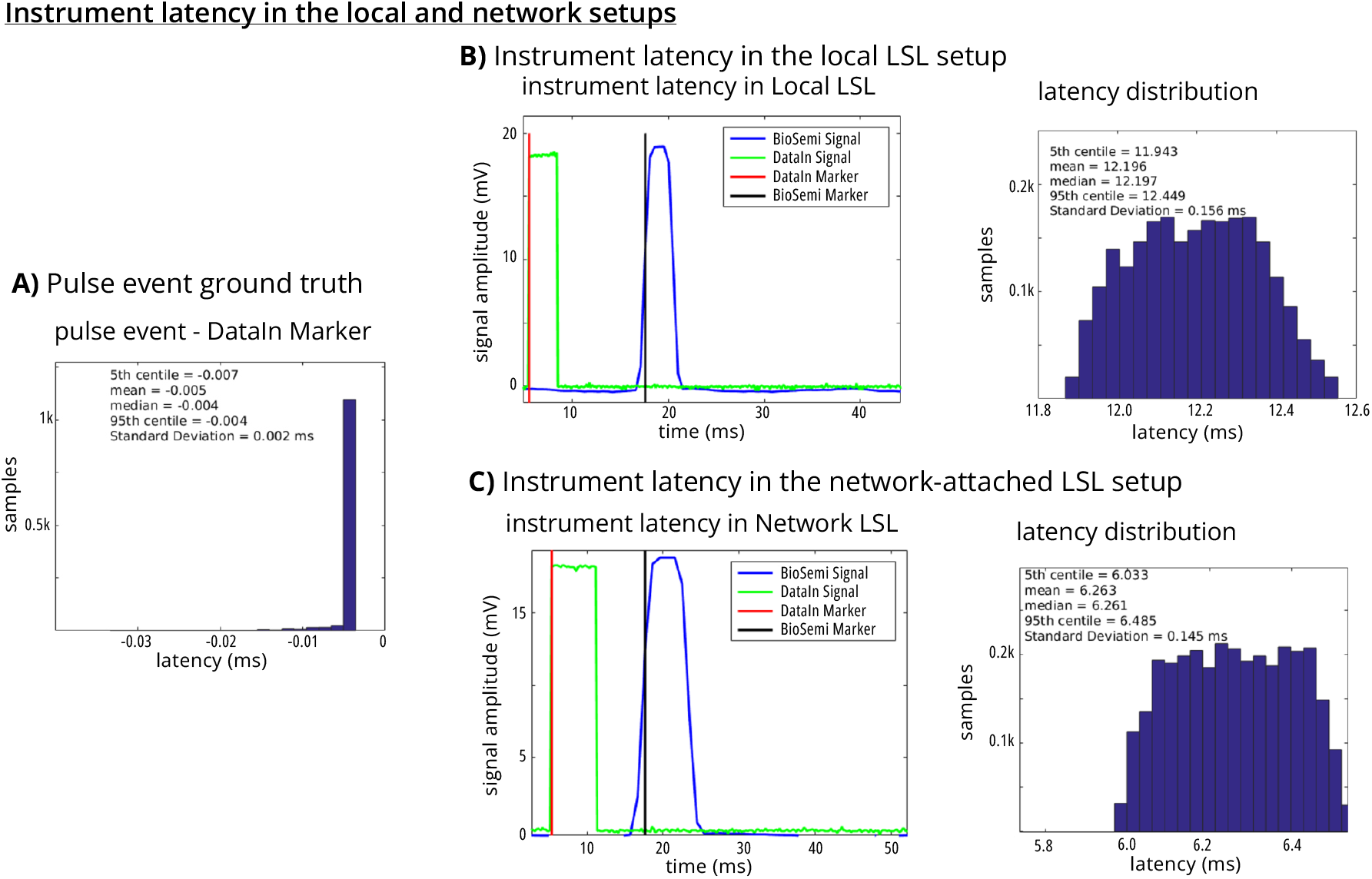
Single-machine (local) and multi-machine synchronization performance. **A**. The Ni-DAQ outputs a *pulse event* to the computer as an LSL inlet everytime a pulse signal is generated. A Ni-DAQ input records the output signal and sends it to another LSL inlet. The *DataIn Marker* is created from this input after as the pulse is detected. **B** *DataIn* and *BioSemi* are recorded the same signal on the same computer. The *DataIn* and *BioSemi* Markers indicate pulse detection by each instrument, respectively. **C** *DataIn* and *BioSemi* are recorded the same signal but on *two* separate machines attached by a wired network. Computationl overhead of recording multiple singals on a local machine may have attributed to the larger offset on the local setup compared to the netwro-attached setup.

### B. Instrument Latency in a Networked LSL Setup

To assess the setup offset of the instrument (in this example the BioSemi amplifier) in a distributed network, we separated the program controlling the NI-Daq (sending the DataIn Marker and storing *pulse events*), the program sending the BioSemi stream, and LabRecorder to network-attached computers. The results showed an even smaller setup offset between the DataIn Marker and the BioSemi Marker than the results observed in the single-machine LSL performance test (here, networked offset: 6.26 ms, vs. local offset: 12.20 ms, (Figure 4C). The offset jitter (presented as the standard deviation of the offset, (Figure 4C) was 145 microseconds, similar to the results from the local network experiment.

This offset decrease might have arisen from the separation, here, of the BioSemi and NI-Daq machines and potentially by faster performance of the BioSemi application and the associated driver running on Windows 10. However, the total setup delay for a given instrument is frequently dominated by device transmission delays, including large on-device buffer sizes that are only periodically transmitted, wireless (e.g., Bluetooth) protocol transmission latencies, and may add up to several 10s of milliseconds. Such discrepancies underpin the importance of testing setup offset (including device throughput) for all devices and configurations before recording experiment data. Setup offsets can be manually added to the metadata while the other potential ad-hoc offsets caused by the network delay or asynchrony would be recorded into the XDF automatically. Both types of offsets will be addressed upon importing the XDF files with the help of the LSL Time synchronization protocol (II-B5) and using the load_xdf function (https://github.com/xdf-modules/xdf-Matlab/blob/master/load_xdf.m).

### C. Multi-instrument Synchronization

To explore the synchronization performance of multimodal recordings on a single PC, a typical research use-case scenario, we measured the jitter between professional-grade acquisition devices (Noraxon Ultium EMG combined with a Labjack T7-pro, and Ant Neuro EEG) as well as a consumer-grade webcam (Logitech C920S HD Pro Webcam) using a standard laptop (Lenovo X1 Carbon Gen 7). To avoid potential delays due to hardware-related TTL triggering, we created two synchronized square wave analog signals, appropriately scaled and conditioned according to device specifications, and injected them directly into EEG and EMG electrodes respectively. These bipolar signals were then acquired and streamed over the local network as physiological data. We simultaneously generated a blinking LED via an Arduino Zero device, captured it via the webcam, and streamed it via LSL over the network along with the EEG and EMG data (Figure 5A).

**Fig. 5.**
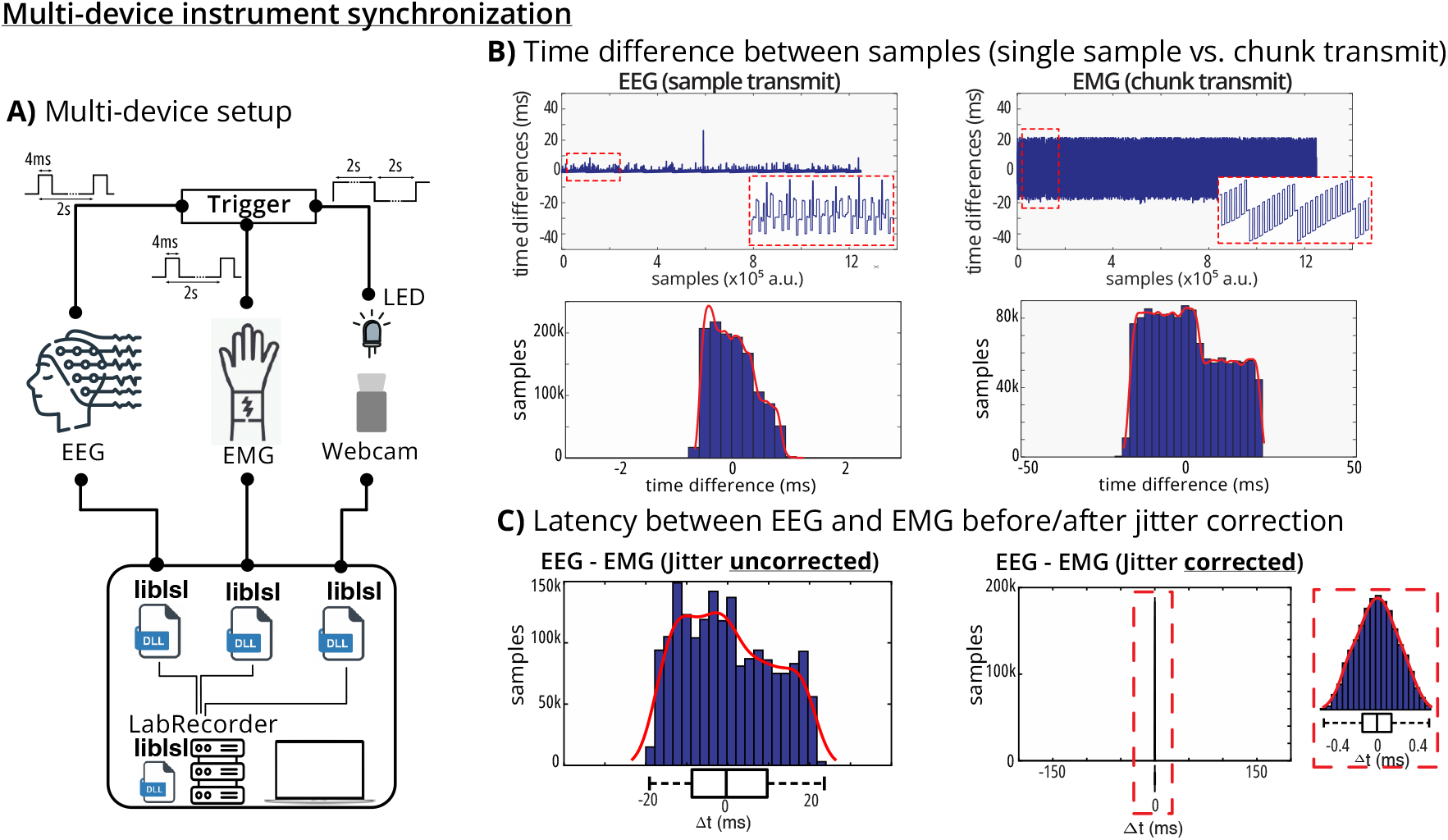
Multi-device synchronization on a local machine. **A**. The trigger signal was recorded by an EEG, EMG and a webcam recording device. Each device transmitted their recording to a single machine. Red dashed boxes indicate zoomed-in looks to the data. **B** EEG signal was tranmistted at each sample, while EMG signal was transmitted by about 20ms chunks. **C** After correcting for the signal offset and jitter, the difference between the EEG and EMG signals was <0.5ms.

The experimental setup consisted of three data streams: EEG sampled at approximately 2000 Hz, EMG sampled at similar rates, and webcam data captured at standard 30 frames per second. All streams were recorded using LabRecorder software in XDF format for 10 minutes. Data were analyzed using MATLAB (R2024b) and imported using both default parameters (HandleJitter = true) and with jitter handling disabled (HandleJitter = false). The timing of rising fronts of the EEG and EMG square waves and LED activation times were computed, subtracted pairwise, and centered to the mean to create Camera vs. EEG, Camera vs. EMG, and EEG vs. EMG jitter distributions.

We observed distinct transmission characteristics between different device types (Figure 5B). The EEG device demonstrated single-sample transmission with minimal jitter, while the EMG device used chunk-based transmission resulting in characteristic periodic timing patterns. The webcam showed more irregular timing behavior typical of consumer-grade devices with variable frame rates Figure 6A).

**Fig. 6.**
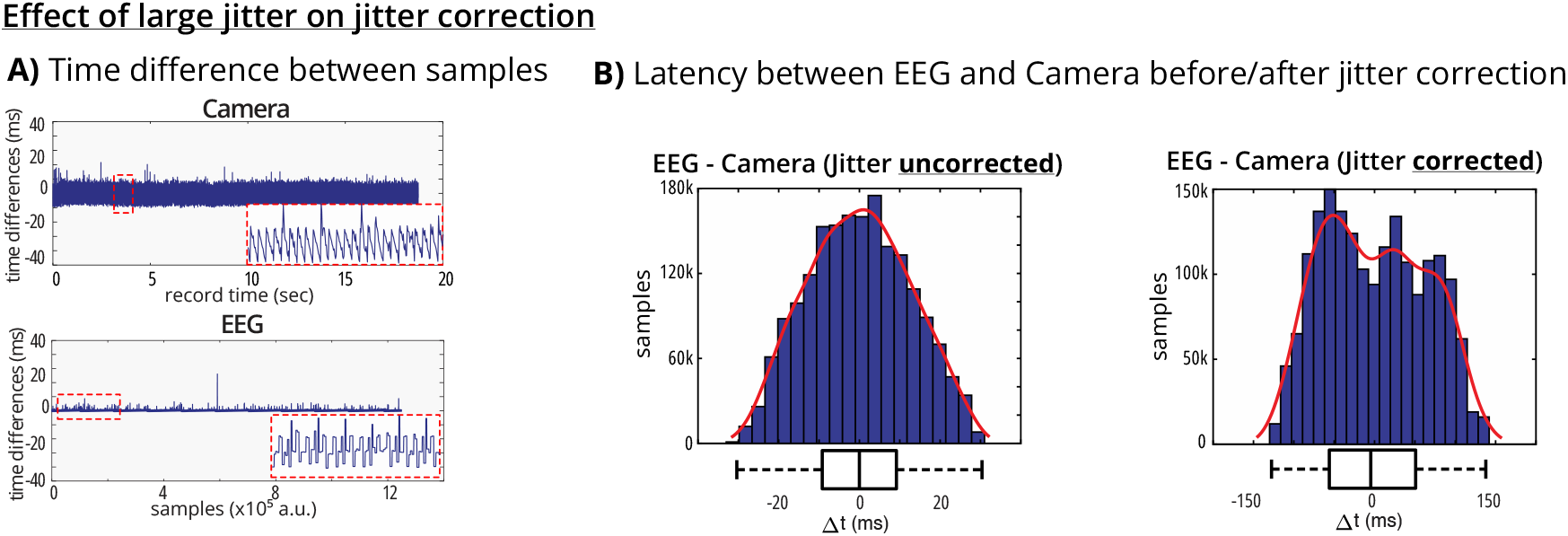
Effect of jitter correction for large jitter consumer-grade instruments. **A)** Time difference between samples for camera (top) and EEG (bottom) streams, showing irregular timing behavior for consumer-grade devices versus professional-grade equipment. Red dashed boxes indicate zoomed-in looks to the data. **B)** Latency distributions between EEG and camera streams before (left) and after (right) jitter correction. Unlike professional-grade device pairs, consumer-grade cameras may show better synchronization with jitter correction disabled, as irregular sampling rates violate Gaussian delay assumptions of the correction algorithm.

The synchronization analysis revealed that sub-millisecond jitter is achievable on standard consumer-grade laptops using default LSL parameters (HandleJitter = true) when professional hardware with “uniform” sampling rates is employed (Figure 5). This was demonstrated for both EEG and EMG devices tested. The jitter-corrected latency between EEG and EMG streams showed a tight distribution centered around zero with standard deviation of approximately 0.5 ms, indicating excellent synchronization performance.

However, synchronization performance varied significantly with device type and parameter settings. Disabling jitter handling (HandleJitter = false) increased jitter by at least one order of magnitude for professional-grade devices, as shown in the EEG-EMG comparison where the uncorrected jitter distribution was substantially broader. Interestingly, for consumer-grade hardware such as a webcam, disabling jitter handling sometimes improved synchronization. This occurs because highly irregular sampling rates violate the Gaussian delay distribution assumptions underlying the jitter correction algorithm (Figure 6).

Our results demonstrate that LSL’s built-in jitter correction is highly effective for professional-grade devices with consistent sampling rates, achieving sub-millisecond synchronization accuracy. However, users should carefully evaluate their specific hardware configurations, as delays between streams can vary over time and differ between hardware setups. Factors such as varying CPU clock speeds due to thermal throttling, operating system prioritization due to workload changes, and hardware-level energy saving features can all affect jitter and delays. Therefore, users are encouraged to test their hardware configurations before critical acquisitions and optimize data analysis pipelines according to their setup’s characteristics.

## IV. PITFALLS AND TWEAKS

LSL’s timing can be influenced by network congestion, device-specific buffering, and clock-drift between hosts. Also, LSL cannot account for internal hardware delays and researchers must determine this delay at least once every time their setup configuration (including adding or removing instruments or netwrok clients, updating drivers or operating system) changes. This section gathers known challenges and hands-on remedies so that researchers can (i) anticipate sources of error before data collection and (ii) apply configuration tweaks or offline corrections to retain sub-millisecond alignment.

### A. Transmission Latency

Transmitting the timestamped data through the LSL network also poses some latency between the outlet and inlets. It is important to reemphasize that data is timestamped by the outlet immediately upon receipt from the data source (e.g., device), and therefore, data transmission latency over the network generally does *not* introduce errors in timestamps. However, such delays may pose some challenges for real-time applications, which want to responsd to received data in a timely manner.

### B. Determining the Setup Offset

As we demonstrated above, adjusting recording times for setup offset is imperative for successful multimodal data acquisition and synchronization. Modifying the setup configuration (e.g., moving an *outlet* from one machine to another) may change the setup offset. Any change in network configurations or updates to their software, drivers, or operating systems should prompt a recheck. Here, we present a simple yet effective procedure to determine setup offset for every instrument, a process similar to that described above in III-A. To determine the setup offset of an instrument, we suggest using a microcontroller unit (e.g., an Arduino) board to send TTL pulses to both the LSL network and to the instrument as a data input (Figure 7). Publishing the TTL pulse as a DataIn Marker can be accomplished through a control software that registers the TTL pulses, or can be directly published by the MCU, since the LSL developer community has provided support for running *liblsl* on some MCUs. The data from the instrument should then be streamed to the LSL network. Both the DataIn Marker and the instrument data should be recorded using *LabRecorder* or an equivalent recording software. The setup should be chosen in a way that most exactly represents the experiment configuration. After reconstructing a marker that corresponds to the TTL pulses from the instrument data (instrument marker, similar to the BioSemi Marker in III-A), the average offset between timestamps of the DataIn markers and the instrument marker is the setup offset. We should note that setup offset can be either positive or negative.

**Fig. 7.**
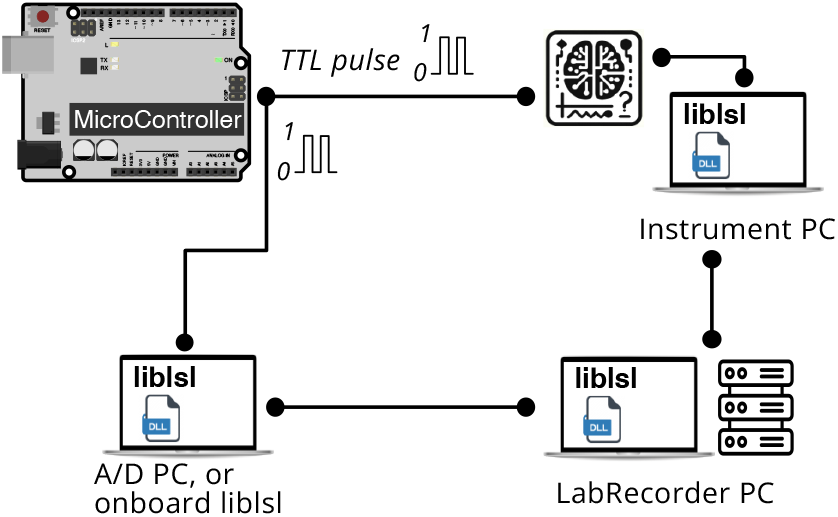
Setup offset determination algorithm. The setup offset can be determined by sending a TTL pulse from a microcontroller board to the LSL network and to the instrument. The instrument data would be streamed to LSL, and the LSL marker would be recorded using LabRecorder. The setup offset would be the average offset between the DataIn marker and the instrument marker.

A positive offset means that the instrument marker occurs after the DataIn marker, indicating an instrument lag in capturing and transmitting the data to the recorder. A negative offset means the instrument marker occurs *before* the DataIn marker; this may happen for sensory triggers (e.g., auditory pulses) where the instrument marker is the time that the trigger pulse is sent to the auditory transducer (e.g., a loudspeaker), while the DataIn marker indicates the time at which the transducer actually produces the pulse.

A successful setup with sub-millisecond internal delay using an affordable MCU board (Arduino) has been benchmarked and could be easily replicated from [Appelhoff and Stenner, 2021]. A commercial solution using dedicated hardware for determining setup offsets is also available from Neurobehavioral Systems, Inc. We again strongly encourage researchers to use these instruments to determine the setup offset and also to verify LSL’s determination of network delays.

### C. Common Device and Network Issues

LSL can address some known hardware failures or network connectivity issues. Sometimes, a hardware device may exhibit a significant change in sampling rate (e.g., in our experience, a webcam that frequently switches between 30 and 60 frames per second) or suffer from high and variable packet loss (e.g., a Bluetooth device that goes in and out of operational range). In these cases, the load_xdf’s attempt to linearly smooth the timestamps will significantly (even catastrophically) distort the data. This can be checked by comparing the effective sampling rate as quantified by load_xdf (as the number of samples divided by the recording length) with the sampling rate reported in the device metadata. If these two sampling rates are not close to each other, we suggest calling load_xdf with the flag ‘HandleJitterRemoval’ set to false. Oftentimes it is possible to recover such recordings with some manual effort thanks to XDF’s policy to record all underlying ground-truth timing data. A similar issue can arise by using LSL through a wireless local area network (WLAN). If there are multiple streams on a heavily utilized WLAN, the clock offset packet exchange can sometimes overload the network and cause gaps in the data. In this case, it may be appropriate to optimize the LSL configuration file for WLAN. The recommended settings for WLANs are:

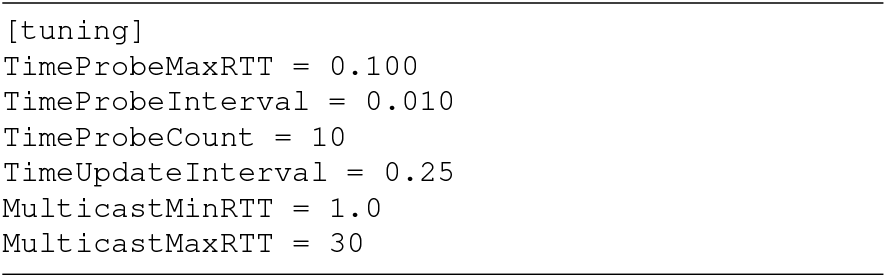

This text can be placed in a file called lsl_api.cfg. If this file is in the same folder as the device’s LSL application, these settings would only be applied to the device. If the file is in ∼/lsl_api/, the changes would be applied to the user globally. If the file is placed in an /etc folder (C:\etc on Windows), the tweaks will be global for all users.

Since applications can supply their own time stamps upon submitting a sample to LSL, potentially outside of the control of the user, it is possible to selectively ignore such time stamps via the user-facing configuration file. This can be necessary when a third-party application uses non-standard time stamps (e.g., from an alternative clock source such as on-device clocks). Since LSL tracks time offset between host machines and not between arbitrary application-chosen clocks, in such cases the recorded data would appear mutually unsynchronized. To rectify this, the user can put the following lines into their `lsl_api.cfg`:

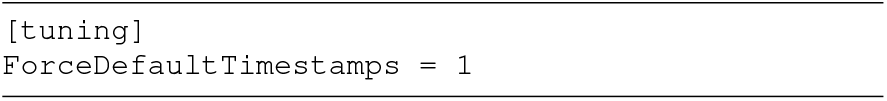

### D. Use in Neurostimulation

A natural extension of LSL’s capabilities is its integration with stimulation paradigms such as transcranial magnetic stimulation (TMS), transcranial electrical stimulation (tES), transcranial ultra-sound stimulation (TUS) and others. LSL can facilitate such setups by recording stimulation onset event markers or the stimulus trains themselves at their native resolution, which allows for post-hoc correlation analysis with respect to neural data. LSL’s ability to access neural data with low transmission delay also facilitates time-synchronized paradigms, including phase-locked or neural burst triggered neurostimulation [Shirinpour et al., 2020].

Integration approaches generally fall into hardware-based solutions (manufacturer-integrated systems^9^ or third-party bridges) and software-based coordination through frameworks, and both solutions can benefit from impleming LSL as their biosignal and trigger sycnhronization framework. Recent comparative studies demonstrate that while hardware-based synchronization achieves superior timing precision, software-based LSL approaches offer greater experimental flexibility for multi-device integration [Miziara et al., 2025]. Advanced closed-loop systems now achieve sub-millisecond precision through novel synchronization methods [Kahilakoski et al., 2025], indicating the field’s rapid evolution toward sophisticated real-time paradigms. When implementing such paradigms, it is important to assess timing requirements and measure both timing error and transmission latency of the envisioned LSL setup. When participant safety considerations arise from timing imperfections (e.g., network latency spikes from wireless connections), researchers should consider acquiring data directly from hardware to drive closed-loop stimulus generation, avoiding network links along the signal path. LSL can facilitate development of such tailored setups through its broad suite of open-source device integrations, which can be repurposed to build direct data paths with minimum latency. Since many devices allow only single-client access, the same program can optionally generate LSL streams for recording purposes, as submitting data to LSL outlets is non-blocking and completes within microseconds with low jitter.

We view this as an important area for future development, and invite and encourage collaboration with researchers working on concurrent stimulation-recording setups to extend LSL’s utility and safety in neuromodulation research.

## V. SUMMARY AND CONCLUSION

The Lab Streaming Layer is a now well-established, reliable and easy-to-use multimodal signal acquisition, transmission, and recording platform tuned for synchronously recording multimodal brain and behavioral data. Oftentimes, using LSL with a given device can be as simple as enabling LSL support in a vendor-provided data acquisition software, if supported, or using one of the existing open-source integrations for the device, and recording the data on the same or another machine with the *LabRecorder* or another LSL-compatible recording tool. However, LSL also scales to complex setups involving multiple machines and several dozen acquisition devices or data streams. In one multiperson, multiple touchscreen simulation [Kothe et al., 2018], we successfully used LSL to record from over 40 LSL data streams^10^ in recording sessions lasting multiple hours.

Recent benchmarks have demonstrated that LSL achieves sub-millisecond synchronization accuracy [Blum et al., 2021], [Chuang et al., 2021], [Iwama et al., 2022], which is on par with or surpasses the timing precision of most existing software-based multimodal acquisition frameworks used in neuroscience. For example, BRAND reported up to 0.5ms in its “inter-node” communication, that is prior to running additional processing or feature extraction pipelines [Ali et al., 2023]. The Falcon framework also reported <1ms latency for Neuralynx hardware, but identified that the latency can increase to multiple milliseconds for long recordings [Ciliberti and Kloosterman, 2017]. Since LSL periodically quantifies the clock offsets and round-trip times between streams, its synchronization accuracy is not affected with the recording length. Our exemplar tests support the excellent sub-millisecond accuracy of the LSL timestamps. As our tests also showed, distributing the computational load of processing multiple streams across separate network-attached machines can at times outperform the setup offset (and latency) achieved by capturing all data streams on a single, perhaps heavily loaded, machine, which is made trivial thanks to LSL’s ability to seamlessly discover streams across the network without additional configuration. For users requiring hardware-level synchronization or TTL integration, the commercial LabStreamer device from Neurobehavioral Systems (See III.D) provides a dedicated plug-and-play solution tightly integrated with LSL.

LSL as a purely software-based approach has an inherent limitation when no hardware triggering mechanisms are used, which is that LSL as a network is not aware of any latency occurring *within* the acquisition device or in the device drivers before data reaches the LSL application for the device. While LSL integrations can make reasonable assumptions, and some do, any residual offset in this latency, which typically amounts to a few 10s of milliseconds, should be ascertained prior to conducting a study, ideally through testing using the actual devices and parameter settings to be used during subsequent recordings. A similar limitation applies to event marker time stamps pertaining to button presses or on-screen presentation, where, again, it is recommended to measure the input and/or display latency using off-the-shelf tools such as photodiodes or high frame rate cameras. Lastly, when the consistency of the device sampling rate itself and/or the stability of its setup offset cannot be trusted, it may be necessary to implement a hardware-based data timing device to monitor the process at least for the affected device(s). Therefore, while LSL can recover lost connections and compensate for offsets and jitter, an appropriate initial setup of the instruments and measuring setup offset are imperative for an optimally synchronized multimodal recording.

While LSL accommodates a relatively large buffer to minimize data loss in case of a connection drop or subpar network speed, given a long enough (e.g., a few minutes) network disconnection, the buffer may eventually run out with the resulting loss of data. Similarly, LSL data throughput is limited by network and computer capacity. While many data streams can be easily transferred at multiple KHz rates, some data streams, such as high-definition video, may saturate the bandwidth. In such a case, using lightweight compression before broadcasting the stream or storing the timestamped data on the local machine and only streaming the timestamps through LSL may resolve this issue.

A large ecosystem, transparent codebase and development, zero-configuration, excellent latency management, and reliability have made LSL a go-to solution for synchronized multimodal quantification of brain and behavior. Since its introduction in 2012, LSL has been cited over 2,300 times, with citations accelerating in recent years, reflecting its growing adoption across the scientific community. Researchers can enjoy LSL with minimal and one-time initial setup and be sure that LSL will stream and store their multimodal data streams accurately and reliably. Finally, LSL development thrives on an open and welcoming community of enthusiasts. Anyone can join this effort via LSL’s community hubs.

## CODE AVAILABILITY

The Lab Streaming Layer (LSL) is free, open-source software maintained by dedicated volunteers. The core library and related packages are available at https://github.com/labstreaminglayer, with the core repositories available from the Swartz Center for Computational Neuroscience (SCCN) GitHub: meta-package and core library. Additional resources, documentation, and a list of compatible devices can be found at https://labstreaminglayer.org. The Extensible Data Format (XDF), used for storing LSL data, is also freely available, with tools and specifications accessible at https://github.com/sccn/xdf.

## AUTHOR CONTRIBUTION

CK and SM conceptualized the LSL ecosystem. CK, TM, and SM devised the methodology. CK, TM, and MG worked in-house on the software. CK, TS, DM, CB, TM, and AD worked on and maintained the software as an open-source project. SYS, CK, TS, DM, CB, and TM wrote the manuscript. SYS generated the final visualizations. All authors reviewed and edited the manuscript. SM provided the funding.

## ACKNOWLEDGMENTS

The first version of the LSL software was written at the Swartz Center for Computational Neuroscience, UCSD, funded by the Army Research Laboratory under Cooperative Agreement Number W911NF-10-2-0022 as well as NINDS grant R01NS047293, and by a gift to UCSD from The Swartz Foundation (Old Field, NY).

## Footnotes

1 https://open-ephys.org

2 https://imotions.com/products/imotions-lab/developers/lsl-support/

3 https://pressrelease.brainproducts.com/lsl-viewer/

4 https://zeromq.org

5 https://mqtt.org

6 https://redis.io

7 https://brainflow.org

8 https://www.w3.org/TR/1999/REC-xpath-19991116/

9 e.g., Magstim-EGI integrated EEG+TMS systems or MxN-Pro featuring LSL

10 Two concurrent subjects, each with instruments including a 267-channel BioSemi, microphone, force plate, eye-tracking, three cameras, motion capture, and event marker streams.

## REFERENCES

[Ali et al., 2023] Ali, Y. H., Bodkin, K., Rigotti-Thompson, M., Patel, K., Card, N. S., Bhaduri, B., Nason-Tomaszewski, S. R., Mifsud, D. M., Hou, X., Nicolas, C., Allcroft, S., Hochberg, L. R., Yong, N. A., Stavisky, S. D., Miller, L. E., Brandman, D. M., and Pandarinath, C. (2023). BRAND: A platform for closed-loop experiments with deep network models. bioRxiv, page 2023.08.08.552473.

[ANSI, 2017] ANSI (2017). Standard for consumer EEG file format.

[Appelhoff and Stenner, 2021] Appelhoff, S. and Stenner, T. (2021). In COM we trust: Feasibility of USB-based event marking. Behav. Res. Methods, 53(6):2450–2455.

[Artoni et al., 2017] Artoni, F., Barsotti, A., Guanziroli, E., Micera, S., Landi, A., and Molteni, F. (2017). Effective synchronization of EEG and EMG for mobile brain/body imaging in clinical settings. Front. Hum. Neurosci., 11:652.

[Bannach et al., 2009] Bannach, D., Amft, O., and Lukowicz, P. (2009). Automatic event-based synchronization of multimodal data streams from wearable and ambient sensors. In Lecture Notes in Computer Science, volume 135 of Lecture notes in computer science, pages 135–148. Springer Berlin Heidelberg, Berlin, Heidelberg.

[Blum et al., 2021] Blum, S., Hölle, D., Bleichner, M. G., and Debener, S. (2021). Pocketable labs for everyone: Synchronized multi-sensor data streaming and recording on smartphones with the lab streaming layer. Sensors (Basel), 21(23):8135.

[Breitwieser and Eibel, 2011] Breitwieser, C. and Eibel, C. (2011). TiA – documentation of TOBI interface a. arXiv [cs.NI].

[Bustamante et al., 2021] Bustamante, S., Peters, J., Scholkopf, B., Grosse-Wentrup, M., and Jayaram, V. (2021). ArmSym: A virtual human–robot interaction laboratory for assistive robotics. IEEE Trans. Hum. Mach. Syst., 51(6):568–577.

[Cerone et al., 2022] Cerone, G. L., Giangrande, A., Ghislieri, M., Gazzoni, M., Piitulainen, H., and Botter, A. (2022). Design and validation of a wireless body sensor network for integrated EEG and HD-sEMG acquisitions. IEEE Trans. Neural Syst. Rehabil. Eng., 30:61–71.

[Chuang et al., 2021] Chuang, C.-H., Lu, S.-W., Chao, Y.-P., Peng, P.-H., Hsu, H.-C., Hung, C.-C., Chang, C.-L., and Jung, T.-P. (2021). Near-zero phase-lag hyperscanning in a novel wireless EEG system. J. Neural Eng., 18(6).

[Ciliberti and Kloosterman, 2017] Ciliberti, D. and Kloosterman, F. (2017). Falcon: a highly flexible open-source software for closed-loop neuroscience. J. Neural Eng., 14(4):045004.

[Clisson et al., 2019] Clisson, P., Bertrand-Lalo, R., Congedo, M., Victor-Thomas, G., and Chatel-Goldman, J. (2019). Timeflux: an open-source framework for the acquisition and near real-time processing of signal streams. In 8th International Brain-Computer Interface Conference.

[Deering, 1989] Deering, D. S. E. (1989). Host extensions for IP multicasting. RFC 1112.

14. [Delgan, ]Delgan. loguru: Python logging made (stupidly) simple.

[Dold et al., 2023] Dold, M., Pereira, J., Janssen, M., and Tangermann, M. (2023). Project dareplane for closed-loop deep brain stimulation. Brain Stimul., 16(1):319– 320.

[Drepper, 2011] Drepper, U. (2011). How to write shared libraries. Technical report. Available online as a technical report, for example from https://www.cs.dartmouth.edu/sergey/cs108/ABI/UlrichDrepper-How-To-Write-Shared-Libraries.pdf.

17. [Fielding et al., ]Fielding, R. T., Nottingham, M., and Reschke, J. RFC 9110: HTTP semantics. https://www.rfc-editor.org/rfc/rfc9110.html. Accessed: 2023-7-13.

[Gramfort et al., 2013] Gramfort, A., Luessi, M., Larson, E., Engemann, D. A., Strohmeier, D., Brodbeck, C., Goj, R., Jas, M., Brooks, T., Parkkonen, L., and Hämäläinen, M. (2013). MEG and EEG data analysis with MNE-python. Front. Neurosci., 7:267.

[IEEE SA Standards Board, 2018] IEEE SA Standards Board (2018). IEEE standard for information technology–portable operating system interface (POSIX(TM)) base specifications, issue 7.

[IEEE SA Standards Board, 2020] IEEE SA Standards Board (2020). IEEE standard for a precision clock synchronization protocol for networked measurement and control systems.

[Iwama et al., 2022] Iwama, S., Takemi, M., Eguchi, R., Hirose, R., Morishige, M., and others (2022). Two common issues in synchronized multimodal recordings with EEG: Jitter and latency. bioRxiv.

[Kahilakoski et al., 2025] Kahilakoski, O.-P., Alkio, K., Siljamo, O., Valén, K., Laurinoja, J., Haxel, L., Makkonen, M., Mutanen, T. P., Tommila, T., Guidotti, R., Pieramico, G., Ilmoniemi, R. J., and Roine, T. (2025). NeuroSimo: an open-source software for closed-loop EEG-or EMG-guided TMS. bioRxiv, page 2025.04.05.647342.

[Kang and Wallraven, 2023] Kang, T. and Wallraven, C. (2023). Gotta go fast: Measuring input/output latencies of virtual reality 3D engines for cognitive experiments. arXiv [cs.HC].

24. [Kapoulkine, ]Kapoulkine, A. pugixml: Light-weight, simple and fast XML parser for c++ with XPath support.

25. [Kohlhoff, ]Kohlhoff, C. Boost.asio - 1.82.0. https://www.boost.org/doc/libs/1_82_0/ doc/html/boost_asio.html. Accessed: 2023-7-1.

[Koranne, 2011] Koranne, S. (2011). Boost c++ libraries. In Handbook of Open Source Tools, pages 127–143. Springer US, Boston, MA.

[Kothe et al., 2018] Kothe, C., Mullen, T., and Makeig, S. (2018). Strum: A new dataset for neuroergonomics research. In 2018 IEEE International Conference on Systems, Man, and Cybernetics, pages 77–82. IEEE.

[Kothe and Makeig, 2013] Kothe, C. A. and Makeig, S. (2013). BCILAB: a platform for brain-computer interface development. J. Neural Eng., 10(5):056014.

[Levitt et al., 2022] Levitt, J., Yang, Z., Williams, S. D., Lütschg Espinosa, S. E., Garcia-Casal, A., and Lewis, L. D. (2022). EEG-LLAMAS: an open source, low latency, EEG-fMRI neurofeedback platform. bioRxiv.

[Lopes et al., 2015] Lopes, G., Bonacchi, N., FrazÃo, J., Neto, J. P., Atallah, B. V., Soares, S., Moreira, L., Matias, S., Itskov, P. M., Correia, P. A., Medina, R. E., Calcaterra, L., Dreosti, E., Paton, J. J., and Kampff, A. R. (2015). Bonsai: an event-based framework for processing and controlling data streams. Front. Neuroinform., 9.

[Maidhof et al., 2014] Maidhof, C., Kästner, T., and Makkonen, T. (2014). Combining EEG, MIDI, and motion capture techniques for investigating musical performance. Behav. Res. Methods, 46(1):185–195.

[Makeig et al., 2009] Makeig, S., Gramann, K., Jung, T.-P., Sejnowski, T. J., and Poizner, H. (2009). Linking brain, mind and behavior. Int. J. Psychophysiol., 73(2):95–100.

[Martin et al., 2010] Martin, J., Burbank, J., Kasch, W., and Mills, D. L. (2010). RFC 5905: Network time protocol version 4: Protocol and algorithms specification. https://datatracker.ietf.org/doc/rfc5905/. Accessed: 2023-11-27.

[Merino-Monge et al., 2020] Merino-Monge, M., Molina-Cantero, A. J., Castro-Garcia, J. A., and Gomez-Gonzalez, I. M. (2020). An easy-to-use multi-source recording and synchronization software for experimental trials. IEEE Access, 8:200618–200634.

[Miziara et al., 2025] Miziara, I. M., Fallon, N., Marshall, A., and Lakany, H. (2025). A comparative study to assess synchronisation methods for combined simultaneous EEG and TMS acquisition. Sci. Rep., 15(1):12816.

[Möller et al., 2008] Möller, B., Morse, K. L., and Lightner, M. (2008). HLA evolved – a summary of major technical improvements.

[Renard et al., 2010] Renard, Y., Lotte, F., Gibert, G., Congedo, M., Maby, E., Delannoy, V., Bertrand, O., and Lécuyer, A. (2010). OpenViBE: An open-source software platform to design, test, and use brain–computer interfaces in real and virtual environments. Presence (Camb.), 19(1):35–53.

[Santamaría-Vázquez et al., 2023] Santamaría-Vázquez, E., Martínez-Cagigal, V., Marcos-Martínez, D., Rodríguez-González, V., Pérez-Velasco, S., Moreno-Calderón, S., and Hornero, R. (2023). MEDUSA(c): A novel python-based software ecosystem to accelerate brain-computer interface and cognitive neuroscience research. Comput. Methods Programs Biomed., 230(107357):107357.

[Schalk et al., 2004] Schalk, G., McFarland, D. J., Hinterberger, T., Birbaumer, N., and Wolpaw, J. R. (2004). BCI2000: a general-purpose brain-computer interface (BCI) system. IEEE Trans. Biomed. Eng., 51(6):1034–1043.

[Shirinpour et al., 2020] Shirinpour, S., Alekseichuk, I., Mantell, K., and Opitz, A. (2020). Experimental evaluation of methods for real-time EEG phase-specific transcranial magnetic stimulation. J. Neural Eng., 17(4):046002.

[Weber et al., 2021] Weber, D., Hertweck, S., Alwanni, H., Fiederer, L. D. J., Wang, X., Unruh, F., Fischbach, M., Latoschik, M. E., and Ball, T. (2021). A structured approach to test the signal quality of electroencephalography measurements during use of head-mounted displays for virtual reality applications. Front. Neurosci., 15:733673.

